# IgaA protein, GumB, has a global impact on the transcriptome and surface proteome of *Serratia marcescens*

**DOI:** 10.1101/2022.04.27.489824

**Authors:** Nicholas A. Stella, Eric G. Romanowski, Kimberly M. Brothers, Robert M. Q. Shanks

**Author notes:** To whom correspondence should be addressed Robert M. Q. Shanks, EEI 1015, 203 Lothrop Street, Department of Ophthalmology University of Pittsburgh Pittsburgh, PA 15213.

## Abstract

Bacterial stress response signaling systems, like the Rcs system, can be triggered by membrane and cell wall damaging compounds including antibiotics and innate immune system factors. These regulatory systems help bacteria survive envelope stress by altering the transcriptome resulting in protective phenotypic changes that may also the influence the virulence of the bacterium. This study investigated the role of the Rcs stress response system using a clinical keratitis isolate of *S. marcescens* with a mutation in the *gumB* gene. GumB, an IgaA ortholog, inhibits activation of the Rcs system, such that mutants have overactive Rcs signaling. Transcriptomic analysis indicated that approximately 15% of all *S. marcescens* genes were significantly altered with two-fold or greater changes in expression in the Δ*gumB* mutant compared to the wild type indicating a global transcriptional regulatory role for GumB. We further investigated the phenotypic consequences of two classes of genes with altered expression in the Δ*gumB* mutant expected to contribute to infections: serralysin metalloproteases PrtS, SlpB and SlpE, and type I pili coded by *fimABCD*. Secreted fractions from the Δ*gumB* mutant had reduced cytotoxicity to a corneal cell line, and could be complemented by induced expression of *prtS*, but not cytolysin *shlBA*, phospholipase *phlAB*, or flagellar master regulator *flhDC* operons. Proteomic analysis, qRT-PCR, and type I pili dependent yeast agglutination indicated an inhibitory role for the Rcs system in adhesin production. Together these data demonstrate that GumB and the Rcs stress response system control *S. marcescens* virulence factors beyond the ShlA cytolysin.

**IMPORTANCE:** Previous studies indicate that the bacterial Rcs system is a key regulator of envelope stress. This study demonstrated that activation of the Rcs system had a global impact on the transcriptome of a clinical isolate of *S. marcescens* including decreased expression of cytotoxic serralysin metalloproteases and biofilm promoting type I pili. These results give mechanistic insight into how the Rcs system contributes to pathogenesis.

## INTRODUCTION

The Rcs system is a multicomponent signal transduction system common to the order Enterobacterales (1–3). The Rcs signaling system, both during infection and under environmental conditions, modifies the cell surface to promote survival. The main components of the Rcs system include the response regulator RcsB and histidine kinase RcsC and phosphorelay protein RcsD. Additional components include the outer membrane protein RcsF and inner membrane protein IgaA that activate and inhibit the Rcs system respectively, and RcsB binding proteins such as RcsA that modify RscB promoter specificity (Figure 1). This system was originally identified as a regulator of capsular synthesis and is known in several species to regulate expression of genes that protect the bacteria from osmotic changes, antimicrobial peptides such as polymyxin B, and the peptidoglycan degrading enzyme lysozyme (3), even when the bacterium is highly resistant to the damaging agent (4).

**Figure 1.**
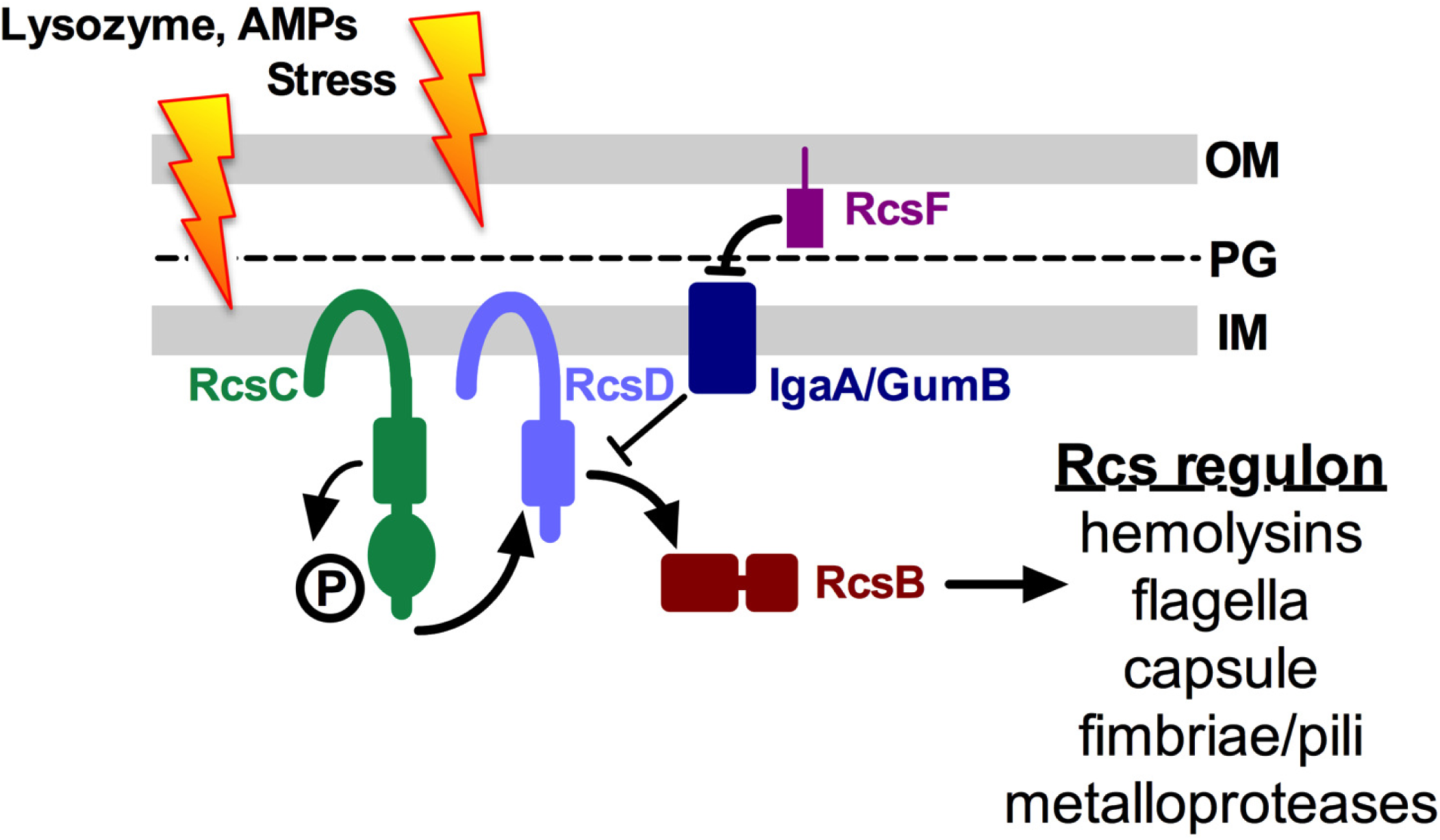
Model for the Res stress response system. This diagram shows the Res phosporelay system that responds to outermembrane (OM) and peptidoglycan cell wall (PG) stresses. This signal transduction system in turn regulates transcription of numerous genes through control of the RcsB transcription factor. Inner membrane protein (IM) GumB (lgaA_SM_) inhibits Res activity during non-stressful conditions, such that mutation of *igaA/gumB* constitutively derepresses RcsB whereas mutation of *rcsB* inactivates the Res system. RcsA (not shown) can bind to and modulate RcsB promoter specificity. In this study we test the hypothesis that the Res system regulates fimbriae and metalloprotease production that in turn inflences the biofilm and cytotoxic potential of the bacteria.

IgaA (intracellular <u>g</u>rowth <u>a</u>ttenuator) family proteins fine tune and inhibit the Rcs system response through interactions with RcsD (5). The first IgaA protein identified was UmoB from *Proteus mirabilis*, found in a screen for flagellar regulators (6). The IgaA protein was later identified in *Salmonella enterica* serovar Typhimurium as point mutations in *igaA* that enabled growth in fibroblast cells (7). Like IgaA in *S. enterica*, the *Escherichia coli* protein, YrfF, is an essential gene (8, 9). In *Serratia* sp ATCC39006 the IgaA protein regulates CRISPR-based immunity to plasmid and phage DNA (10), and in *S. marcescens* the IgaA family protein GumB regulates multiple phenotypes including microbial pathogenesis, biosynthesis of antimicrobial secondary metabolites, capsular polysaccharide, and the *shlA* cytolysin (11–14).

The Rcs system has been highly studied in the Enterobacteriaceae bacteria *E. coli* and *S. enterica*, but is much less described in other bacteria such as the *Yersiniaceae* member *S. marcescens*. A handful of studies in the last 10 years have investigated the Rcs system in a variety of *S. marcescens* strains. Pan *et al* observed that *rcsB* mutants of *S. marcescens* environmental isolate JNB5-1 experience global changes in gene expression including those involved in prodigiosin biosynthesis, motility, acid tolerance, and capsular polysaccharide biosynthesis (15). Pigment regulation by the Rcs system was also shown with a clinical isolate (16). The Garcia Véscovi group demonstrated that that RcsB regulated expression of the *shlBA* toxin operon, flagellar regulators *flhDC* and *fliA*, and mediates autophagy in a rodent cell line (17). The *S. marcescens* Rcs system is activated by perturbation of enterobacterial common antigen synthesis and by high osmolarity (18). The Rcs system was also implicated in the biosynthesis of *S. marcescens* outer membrane vesicles suggesting that it has an important role in envelope homeostasis (19).

The isolate used in this study was from an ocular surface infection, and *S. marcescens* is a leading cause of contact lens associated keratitis (20–22). The ocular surface is unusually paucibacterial for a surface-exposed tissue due in part to the hostile environment presented by innate immune factors in the tear film such as lysozyme and antimicrobial peptides (23–25). Given that during infections throughout the body bacteria are exposed to innate immune activators of the Rcs system, it is likely that the Rcs system plays a role in bacterial pathogenesis. Consistent with this model, an earlier study has shown that dysregulation of the Rcs system through mutation of key genes *gumB*, *rcsB*, and *rcsC* led to highly altered keratitis in a rabbit model, with Rcs-inactivated mutants demonstrating increased virulence and Rcs-activated mutants showing reduced pathogenesis (11). The virulence factors responsible for the altered pathogenesis have not been fully established, although likely involved in differential expression of the *shlBA* cytolysin operon and was suppressed in a Δ*rcsB ΔshlB* double mutant (11). In this study, the role of GumB in global gene expression regulation was investigated in order to better understand how the Rcs system influences microbial pathogenesis.

## RESULTS

### GumB has a global impact on the *S. marcescens* transcriptome

Previous studies demonstrated that IgaA-family protein, GumB, was necessary for cytotoxicity and control of virulence factors including bacterial capsule and ShlA cytolysin (13, 14) and pathogenesis in a rabbit keratitis model (11). To identify other genes influenced by GumB that may influence virulence, RNA-sequencing (RNA-Seq) was performed on RNA harvested from *S. marcescens* wild type and Δ*gumB* mutant grown in LB medium to OD_600_ nm of 1.0. Strains are listed in Table S1. For each genotype, three samples, each the combination of three independent cultures, were evaluated. A total of 722 genes had significant 2-fold or higher differences between wild-type and Δ*gumB* samples, with 411 up-regulated and 311 down-regulated in the Δ*gumB* mutant relative to the wild type (Table S2). Table 1 lists selected genes; among these are genes previously determined by qRT-PCR to have differential expression in the Δ*gumB* mutant compared to the wild type including flagella regulator *flhD*, flagellin gene *fliC*, pigment biosynthetic gene *pigA*, capsular polysaccharide gene *wecA*, and cytolysin operon *shlBA* (13, 14).

**Table 1.**
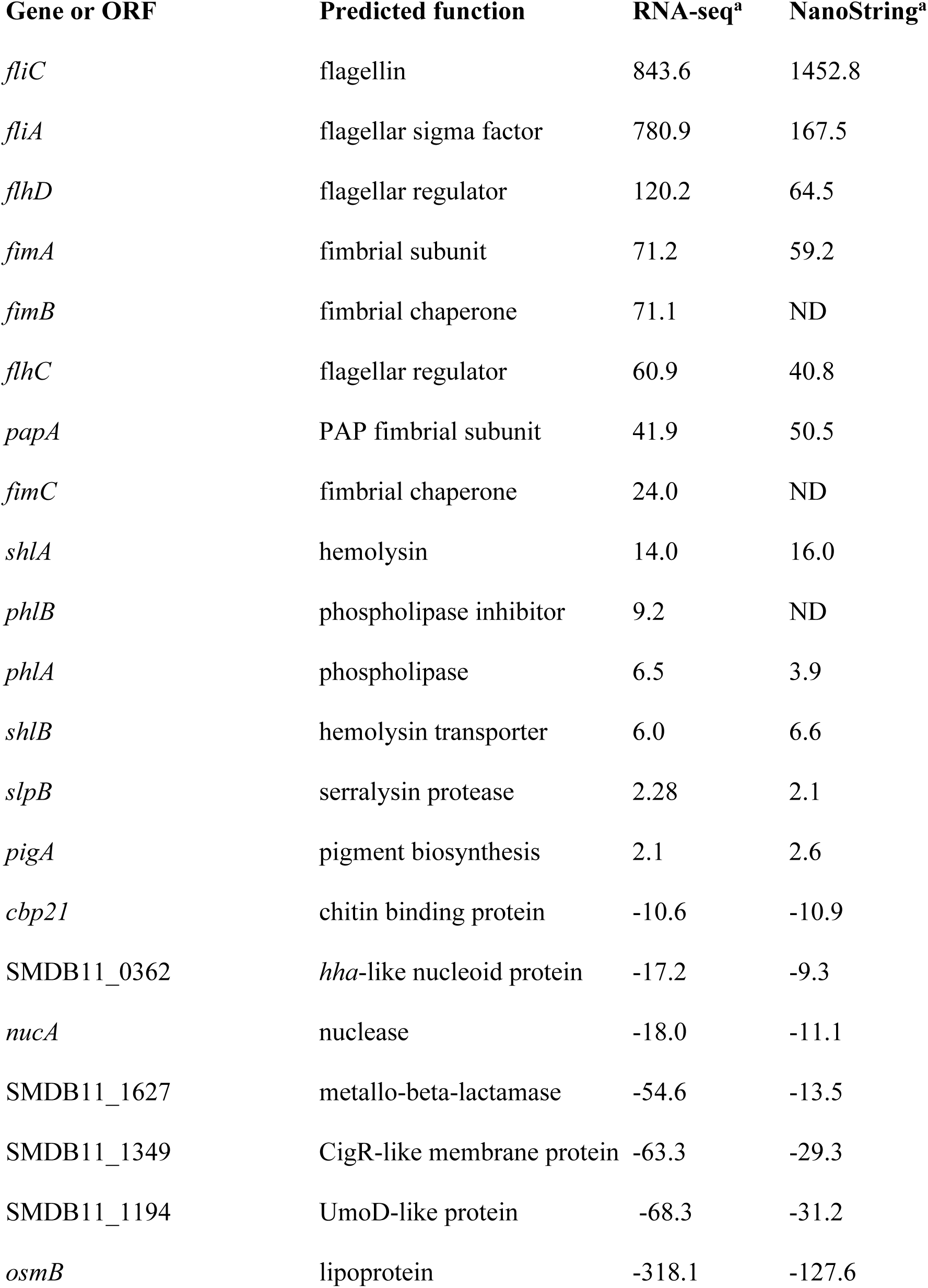
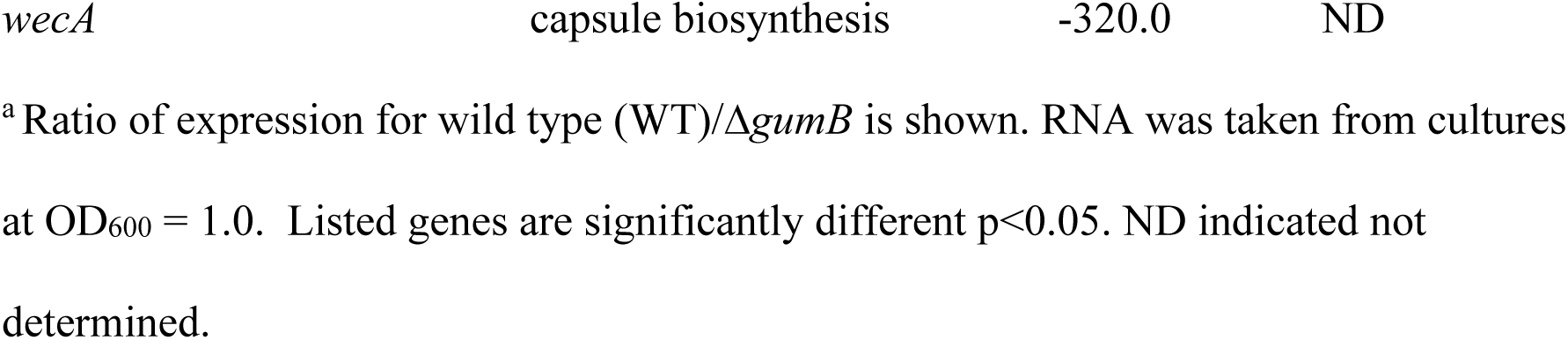
Selected Differentially expressed genes (WT/Δ*gumB*) *in vitro*.

To validate the RNA-Seq expression data, transcripts from a subset of genes were measured using Nanostring technology on independent RNA samples (n=6 for Δ*gumB* and n=8 for wild type). A target set of 70 bacterial genes for Nanostring analysis was developed based on the RNA-Sequencing experiment noted above, orthologs of genes identified in transcriptomic studies of *S. enterica* with reduced function *igaA* alleles (26, 27), and other genes of interest including those that code for potential virulence factors and secondary metabolite biosynthetic proteins. A significant correlation (r=0.9781, p<0.0001 by Spearman analysis) was observed between the RNA-seq and Nanostring analysis of genes with significant differences between the wild type and Δ*gumB* (Figure 2A). Notable results were listed in Table 1. The RNA-sequencing experiment was performed a second time and the results of both RNA-seq experiments correlated significantly (r=0.798, p<0.0001 by Spearman analysis), Figure S1.

**Figure 2.**
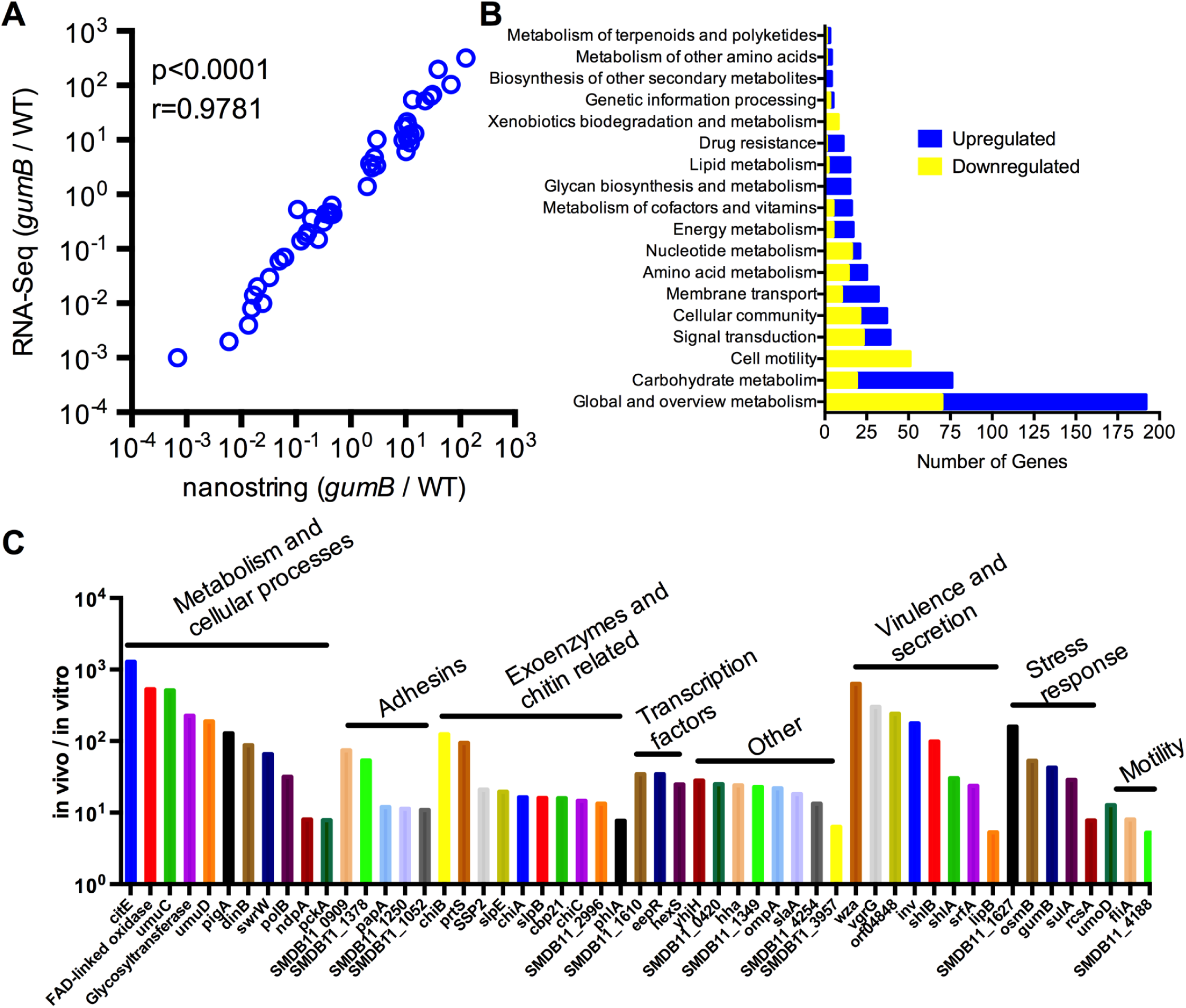
GumB has a global effect on gene expression in *S. marcescens.* **A.** Comparison of RNA-Seq and Nanostring analysis of a subset of genes from RNA isolated from cultures grown to OD_600_= 1 in LB medium. **B.** Genes with significant 2-fold changes were classified according to KEGG pathways. **C.** RNA harvested from corneal infections were compared to RNA from cultures grown to early stationary phase/late log phase in LB medium. Nanostring counts were normalized to ribosomal protein S1p gene expression. Genes with 5-fold or greater differences are shown. *Samples with counts lower than the geometric mean of the negative controls were eliminated.* n=3-5 for *in vivo* and 6-7 for *in vitro* samples. All genes shown were significantly different between the two groups (p<0.05).

KEGG orthology classifications were assigned to differentially expressed genes using GhostKOALA software (28). About 50% of the genes were classified and distributed according to major pathways shown in Figure 2B. More were up-regulated than down-regulated and pathways involved various types of metabolism, cellular motility, signal transduction, and secondary metabolite biosynthesis. Despite reduced expression of several metabolic pathways, previous growth analysis of this Δ*gumB* strain in LB, M9 minimal medium supplemented with glucose, and filtered *Galleria mellonella* homogenate were similar (13, 14). This indicates that the Δ*gumB* mutant is not prototrophic for essential pathways when using glucose as a sole carbon source.

### *S. marcescens* gene expression during microbial keratitis is regulated by GumB

In order to determine *S. marcescens* genes expressed during a relevant infection and whether mutation of *gumB* would have a similar effect on gene expression *in vivo*, RNA was harvested from rabbit corneas 24 hours post injection with ∼1000 CFU and was analyzed using Nano-String. In this infection model, the wild type and mutant are both capable of infecting the eyes and replicating to achieve high bacterial burdens, although the Δ*gumB* mutant induces a notably reduced pathogenic inflammatory response (11).

Genes expressed from the wild-type bacteria *in vivo* are listed in Table S3. The most highly expressed gene *in vivo* was for the RNA binding protein Hfq, which was the second most highly expressed gene from the *in vitro* samples. When bacterial RNA was collected from eyes that had been infected with the *ΔgumB* mutant (n=4) and the wild type (n=5), several differences were observed (Table 2). Of the 70 tested genes five were detected from both groups and differentially expressed (≥2-fold and p<0.05). Two were higher in wild-type samples: *fimA*, a key factor in *S. marcescens* biofilm formation and adherence to mammalian cells (29, 30), and the cytotoxic serralysin family protease gene *slpE* (31). Three genes, detected only in the Δ*gumB* mutant infected eyes, were predicted to be involved in capsule production or were the ortholog of the osmotically induced lipoprotein gene *osmB*. Twenty-four genes were only detected in wild-type infected eyes. These include metalloprotease genes *prtS* and *slpB*, various transcription factor genes *crp*, *flhD*, *oxyR*, *pigP*, and *rcsB,* and toxin/hemolysin genes *shlB* and *shlA*.

**Table 2.**
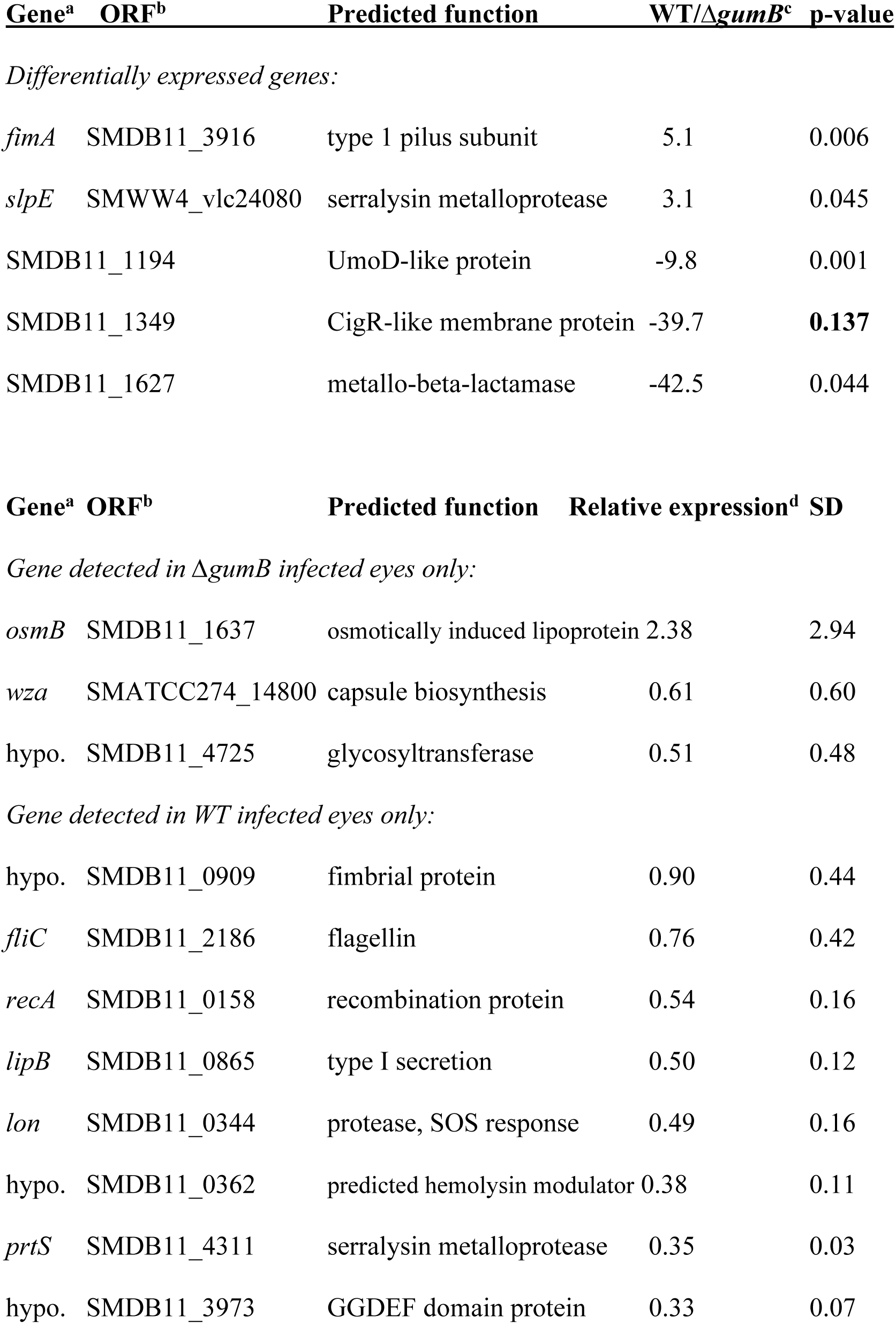
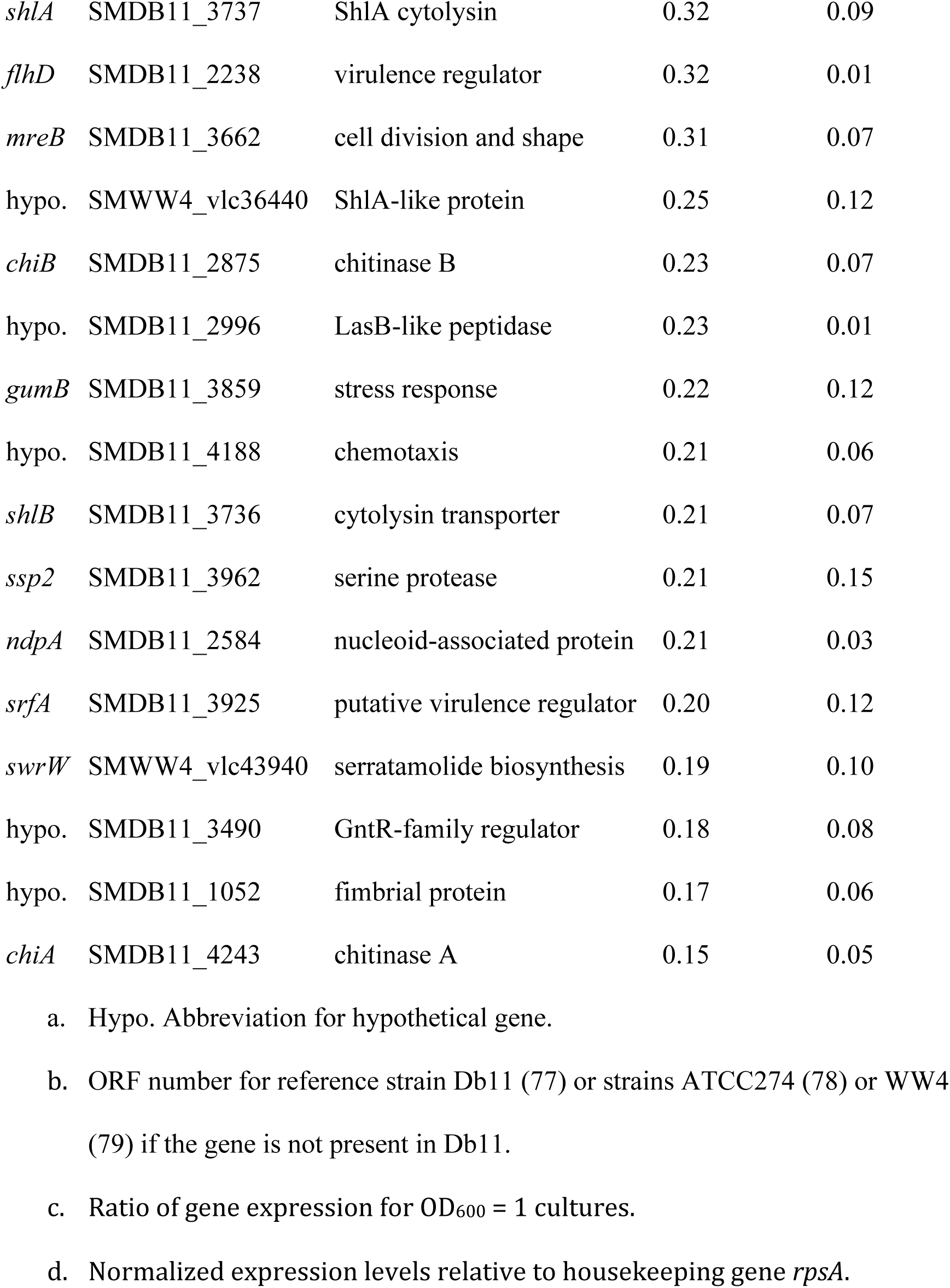
Differentially expressed bacterial genes during keratitis, 24 h post-infection.

| Gene<sup>a</sup> | ORF<sup>b</sup> | Predicted function | WT/$\Delta gumB$<sup>c</sup> | p-value |
| --- | --- | --- | --- | --- |
| *Differentially expressed genes:* |  |  |  |  |
| *fimA* | SMDB11_3916 | type 1 pilus subunit | 5.1 | 0.006 |
| *slpE* | SMWW4_vlc24080 | serralysin metalloprotease | 3.1 | 0.045 |
|  | SMDB11_1194 | UmoD-like protein | -9.8 | 0.001 |
|  | SMDB11_1349 | CigR-like membrane protein | -39.7 | **0.137** |
|  | SMDB11_1627 | metallo-beta-lactamase | -42.5 | 0.044 |

| Gene<sup>a</sup> | ORF<sup>b</sup> | Predicted function | Relative expression<sup>d</sup> | SD |
| --- | --- | --- | --- | --- |
| *Gene detected in $\Delta gumB$ infected eyes only:* |  |  |  |  |
| *osmB* | SMDB11_1637 | osmotically induced lipoprotein | 2.38 | 2.94 |
| *wza* | SMATCC274_14800 | capsule biosynthesis | 0.61 | 0.60 |
| hypo. | SMDB11_4725 | glycosyltransferase | 0.51 | 0.48 |
| *Gene detected in WT infected eyes only:* |  |  |  |  |
| hypo. | SMDB11_0909 | fimbrial protein | 0.90 | 0.44 |
| *fliC* | SMDB11_2186 | flagellin | 0.76 | 0.42 |
| *recA* | SMDB11_0158 | recombination protein | 0.54 | 0.16 |
| *lipB* | SMDB11_0865 | type I secretion | 0.50 | 0.12 |
| *lon* | SMDB11_0344 | protease, SOS response | 0.49 | 0.16 |
| hypo. | SMDB11_0362 | predicted hemolysin modulator | 0.38 | 0.11 |
| *prtS* | SMDB11_4311 | serralysin metalloprotease | 0.35 | 0.03 |
| hypo. | SMDB11_3973 | GGDEF domain protein | 0.33 | 0.07 |

|  |  |  |  |  |  |
| --- | --- | --- | --- | --- | --- |
| 686 | *shlA* | SMDB11_3737 | ShlA cytolysin | 0.32 | 0.09 |
| 687 | *flhD* | SMDB11_2238 | virulence regulator | 0.32 | 0.01 |
| 688 | *mreB* | SMDB11_3662 | cell division and shape | 0.31 | 0.07 |
| 689 | hypo. | SMWW4_vlc36440 | ShlA-like protein | 0.25 | 0.12 |
| 690 | *chiB* | SMDB11_2875 | chitinase B | 0.23 | 0.07 |
| 691 | hypo. | SMDB11_2996 | LasB-like peptidase | 0.23 | 0.01 |
| 692 | *gumB* | SMDB11_3859 | stress response | 0.22 | 0.12 |
| 693 | hypo. | SMDB11_4188 | chemotaxis | 0.21 | 0.06 |
| 694 | *shlB* | SMDB11_3736 | cytolysin transporter | 0.21 | 0.07 |
| 695 | *ssp2* | SMDB11_3962 | serine protease | 0.21 | 0.15 |
| 696 | *ndpA* | SMDB11_2584 | nucleoid-associated protein | 0.21 | 0.03 |
| 697 | *srfA* | SMDB11_3925 | putative virulence regulator | 0.20 | 0.12 |
| 698 | *swrW* | SMWW4_vlc43940 | serratamolide biosynthesis | 0.19 | 0.10 |
| 699 | hypo. | SMDB11_3490 | GntR-family regulator | 0.18 | 0.08 |
| 700 | hypo. | SMDB11_1052 | fimbrial protein | 0.17 | 0.06 |
| 701 | *chiA* | SMDB11_4243 | chitinase A | 0.15 | 0.05 |

a. Hypo. Abbreviation for hypothetical gene.
b. ORF number for reference strain Db11 (77) or strains ATCC274 (78) or WW4 (79) if the gene is not present in Db11.
c. Ratio of gene expression for OD<sub>600</sub> = 1 cultures.
d. Normalized expression levels relative to housekeeping gene *rpsA*.

We compared the *in vivo* and *in vitro* gene expression levels from the Nanostring analysis and observed notable differences (Figure 2C). Classes of genes with significant changes included stress response genes and known and putative virulence factors. Because more bacterial RNA was harvested from the LB cultures, the counts were normalized to the ribosomal S1p protein (*rpsA*) and relative gene expression was compared. The identical trend was observed when counts were normalized by DNA gyrase A gene expression. Most of the 70 tested genes had higher expression or no change *in vivo* compared to *in vitro* growth, notable exceptions were the flagellin gene *fliC*, which was down 3.4-fold, and the type I fimbria gene *fimA*, which had 3.6-fold lower relative expression *in vivo* relative to *in vitro* (p<0.001). Genes with higher relative expression *in vivo* included many known and putative virulence factors including a capsular polysaccharide gene, metalloprotease genes *prtS* (also known as *prtA*), *slpB*, and *slpE*, and the cytolysin operon *shlBA* that had greater than 5-fold higher relative expression in the cornea (Figure 2C). Stress associated genes *hfq*, *oxyR*, and *rcsB* were also elevated greater than 2.5-fold *in vivo* (p<0.01).

### Cytotoxicity of the Δ*gumB* mutant to a human corneal epithelial cell line is decreased in a protease dependent manner

Several genes with reduced expression in the Δ*gumB* mutant from RNA-seq and Nanostring experiments are implicated in cytotoxicity and virulence. These included metalloprotease genes (32), the *shlBA* cytolysin operon (33), the *phlAB* phospholipase operon (34–36), and the *flhDC* transcriptional regulator operon (17, 35, 37) (Table 1). qRT-PCR of RNA harvested from bacteria in late logarithmic and early stationary phase verified these findings (Figure 3A). Similar results were observed with cytotoxic metalloproteases *prtS*, *slpB*, and *slpE* (Figure 3B).

**Figure 3.**
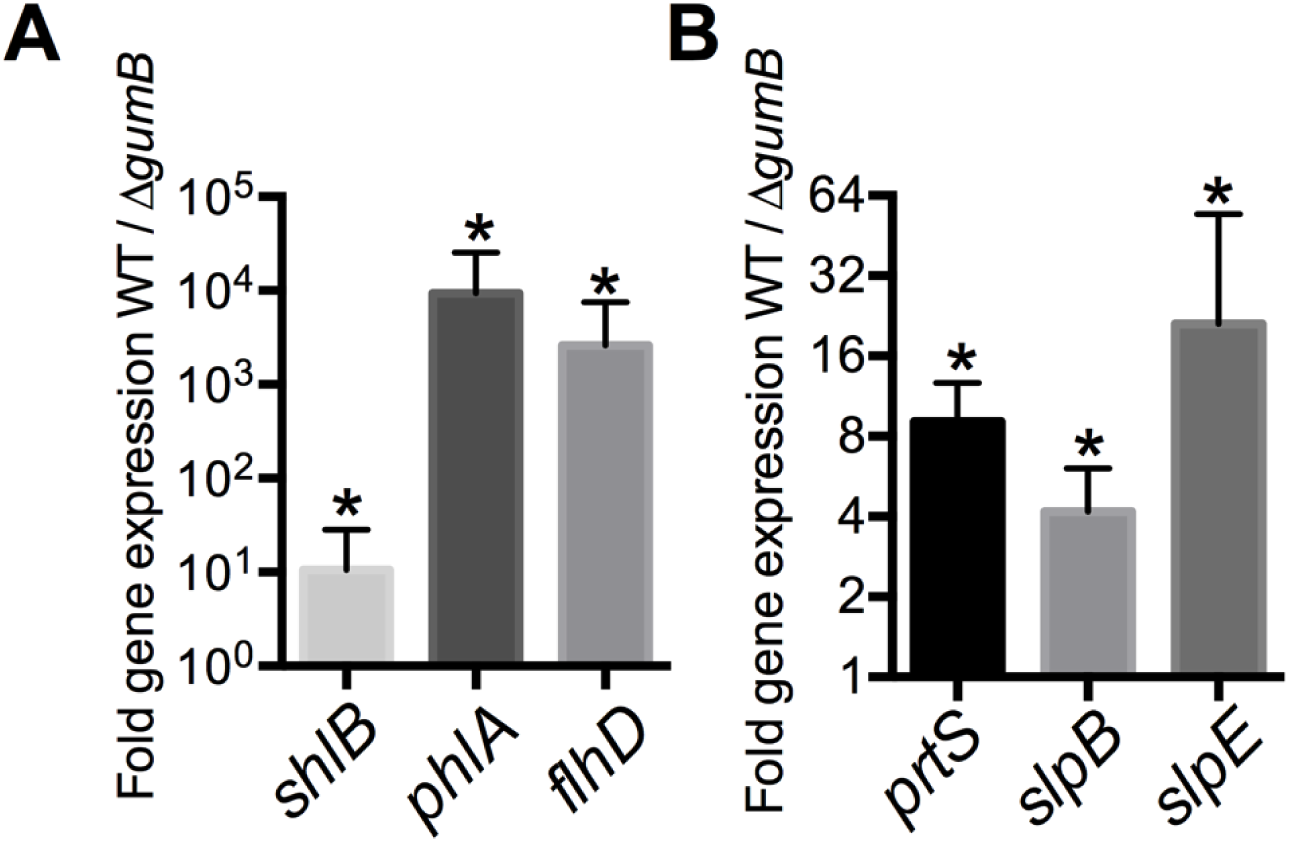
Differential expression of virulence associated genes in the Δ*gumB* mutant. **A-B.** qRT-PCR of gene expression as noted above from cultures harvested at OD_600_ = 1 (or 3 for *ph/A)* in LB medium. Asterisk indicates significant difference between strains, p<0.05. n≥3.

The Δ*gumB* mutant was previously shown to be less cytotoxic to human cell lines when the bacteria were directly incubated with the cell lines and that the reduced cytotoxicity was a consequence of lower ShlA production (13). However, *S. marcescens* is known to have multiple secreted cytotoxic factors, some dependent on bacteria-cell contact and some that are found in bacterial culture supernatants (31, 38, 39). Here we evaluated cytotoxicity of normalized culture supernatants rather than bacterial cells, and observed that the Δ*gumB* mutant was defective in cytotoxicity (Figure 4A) and this could be complemented by wild-type *gumB* on a plasmid when cell viability was assessed using resazurin-based (Figure 4A) and Calcein AM (Figure 4B). This suggested that GumB is necessary for secretion of a cytotoxic compound.

**Figure 4.**
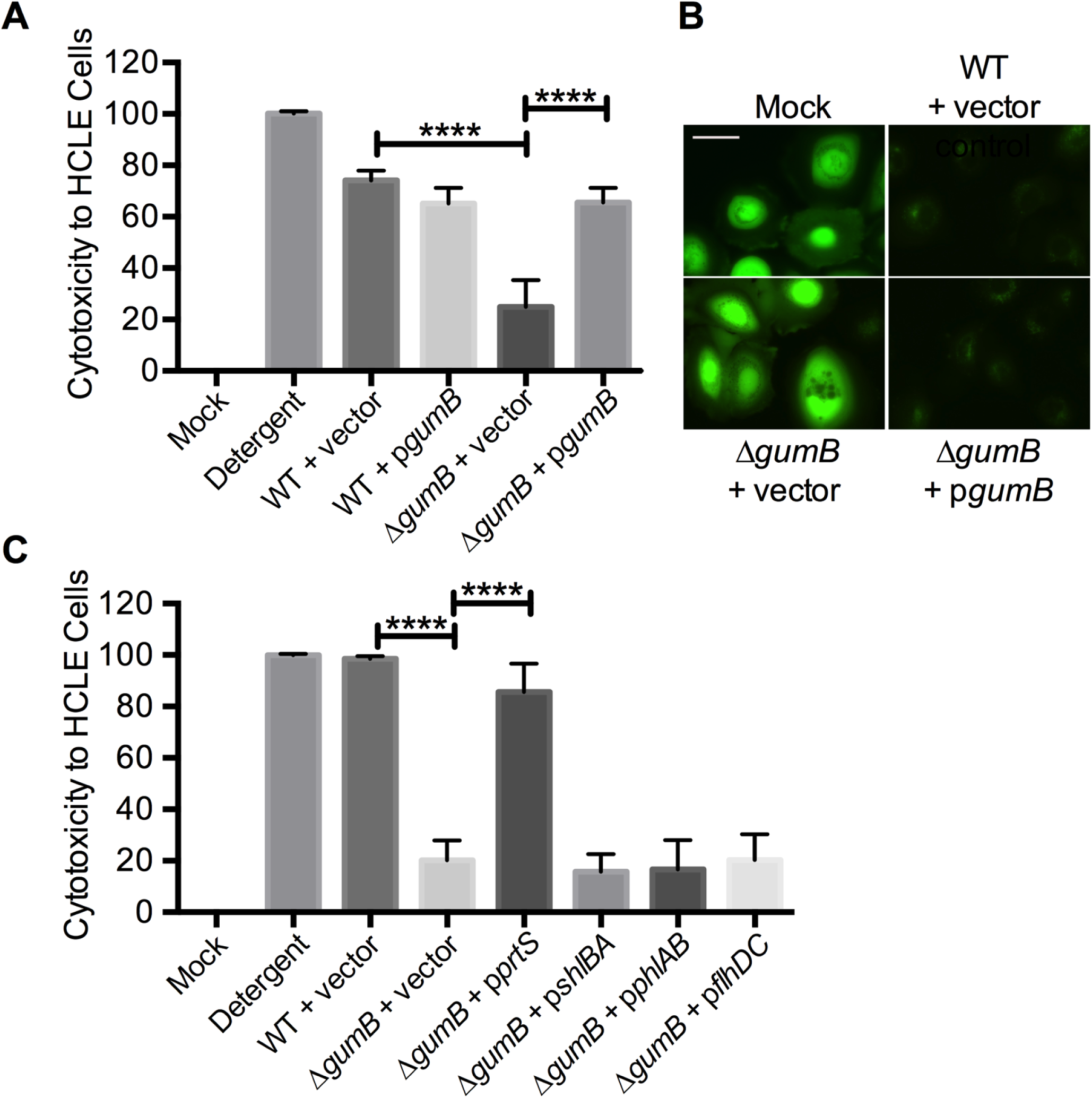
*S. marcescens* cytotoxicity to human corneal cells requires GumB and the Δ*gumB* mutant defect can be reversed through ectopic expression of serralysin (PrtS). **A and C.** Viability of HCLE cells challenged by normalized bacterial supernatants for 4 hours was measured using PrestoBlue®. Detergent (Triton X-100 at 0.25%) and Mock (cell culture growth medium) were set at 100 and 0% cytotoxicity respectively. ≥8, means and SD are shown. Asterisks indicate p<0.0001. Vector control is pMQ132 for panel A and pMQ125 for panel B. **B.** Viability staining of HCLE cells challenged with bacterial normalized supernatants for 4 h and assessed using Calcein AM staining (green) which fluorescently labels live cells.

To determine the secreted cytotoxic factor(s) missing in the Δ*gumB* mutant culture supernatant, we expressed candidate genes that code for cytotoxic factors in the Δ*gumB* mutant and evaluated the cytotoxic capacity of the resulting cells. The candidates were chosen based on corresponding to those genes down-regulated in the Δ*gumB* mutant. This was done using plasmids previously verified to functionally express the cloned gene(s) (13, 35, 39, 40). Supernatants from the Δ*gumB* mutant expressing serralysin gene *prtS*, but not cytolysin operon *shlBA*, or phospholipase operon *phlAB*, restored cytotoxicity to the Δ*gumB* mutant (Figure 4C). A plasmid overexpressing *flhDC*, a positive transcriptional regulator for *shlBA* and *phlAB*, also did not restore cytotoxic potential to the Δ*gumB* mutant (Figure 4C). Together these results suggested that the lack of cytotoxicity of Δ*gumB* mutant secreted factors was due to reduced synthesis of serralysin-family proteases such as PrtS.

### Extracellular protease levels are reduced in the Δ*gumB* mutant

The role of GumB in regulating cytotoxic proteases was analyzed due to reduced transcriptional expression of serralysin family metalloprotease genes (Figure 3B, Table 1). A 2.8-fold reduction in extracellular protease activity was measured from stationary phase Δ*gumB* bacteria using azocasein as a substrate (Figure 5A). The protease defect could be complemented by wild-type *gumB* on a plasmid confirming that GumB is required for full levels of protease activity in the culture supernatants (Figure 5B). These data suggested that the increased Rcs activity in the Δ*gumB* mutant was responsible for the reduced protease production by *S. marcescens*.

**Figure 5.**
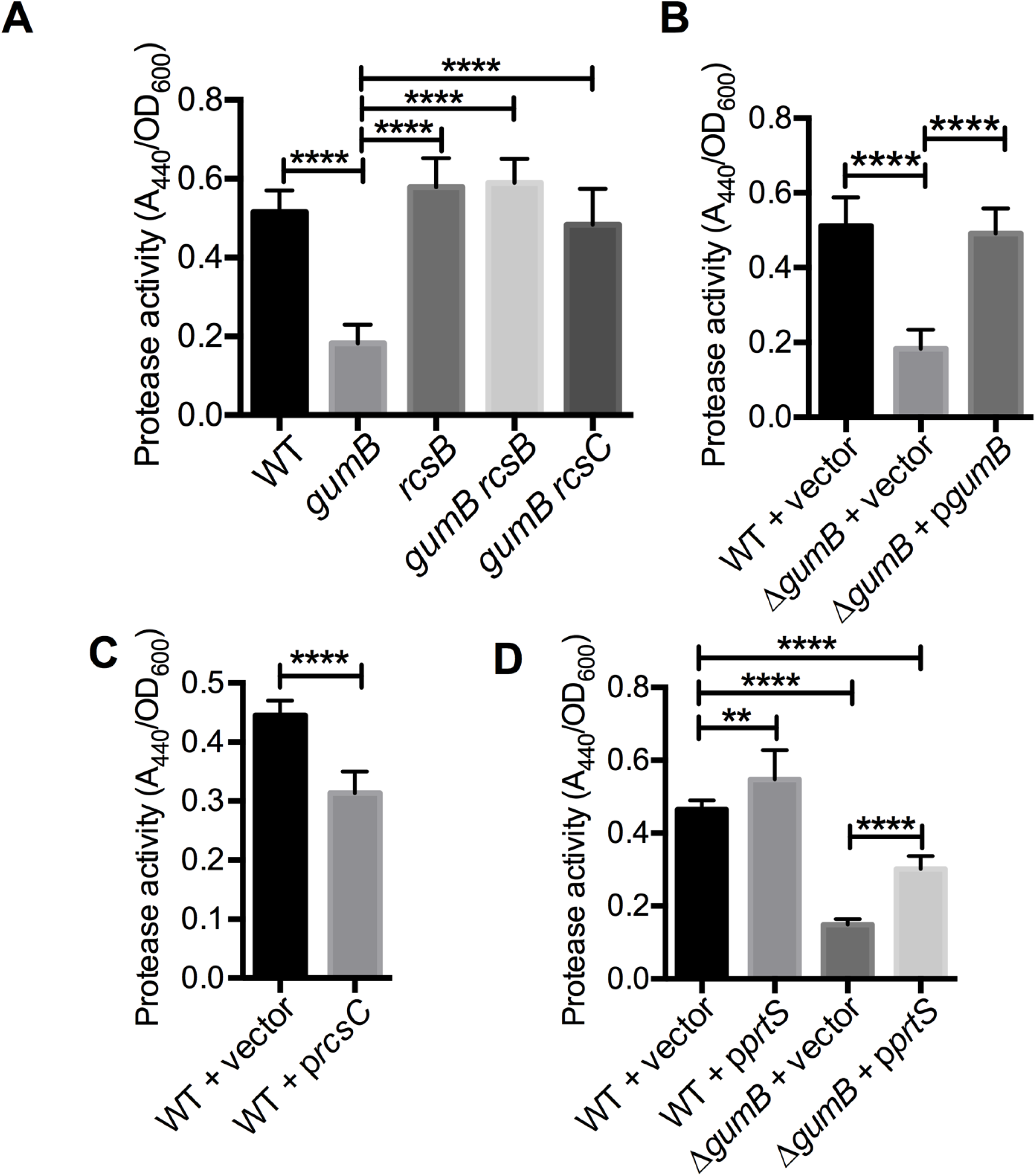
GumB and Res regulation of *S. marcescens* metalloproteases. **A-D.** Protease activity measured using azocasein from stationary phase cultures supernatants normalized to OD_600_= 2. n≥6. ** p<0.01; **** p<0.0001. **D.** The WT + vector (pMQ132) group was significantly different than all other groups (p<0.0001).

To test the model that increased Rcs system activity inhibits protease production in another way, the *rcsC* gene was expressed from a multicopy plasmid in the wild type, which has been shown to enhance Rcs-dependent phenotypes (11, 13). Figure 5C depicts a significant 30% reduction in protease activity measured in the supernatants of the wild type expressing *rcsC* compared to the wild type with the vector control, supporting the model that Rcs activation inhibits *S. marcescens* protease production.

While the expression of the metalloprotease genes were diminished (Figure 3B), it is possible that the Δ*gumB* mutant is defective in secretion of these proteases rather than production. To test this, a representative metalloprotease gene, *prtS*, on a plasmid was expressed from an inducible promoter in the Δ*gumB* mutant. This plasmid partially restored secreted protease activity indicating that the *gumB* mutant is capable of secreting serralysin-family proteases and supporting the model that the secreted protease defect is due to reduced transcription of serralysin proteases rather than the inability to secrete them (Figure 5D).

If the impact of GumB on protease biosynthesis is through over-activation of the Rcs system, then inactivation of Rcs system components downstream of GumB should restore protease activity to the Δ*gumB* mutant. Secreted protease levels were evaluated in Δ*gumB* Δ*rcsB* and Δ*gumB rcsC* double mutants and these were restored to at least wild-type levels (Figure 5A). Mutation of *rcsB* in the wild-type background correlated with a trend toward increased protease activity (12%) that did not achieve significance (p>0.05) (Figure 5A). Together these suggest an inhibitory role for the Rcs system on metalloprotease production.

### Type I pili gene expression and fimbriae-dependent phenotypes are reduced in the Δ*gumB* mutant

Type I pili (fimbriae) are major virulence determinants for many bacteria that promote adhesion to surfaces, such as the cells that line the urinary tract (41). The *fimABCD* operon of different *S. marcescens* strains was demonstrated to be required for attachment to corneal epithelial cells and to a variety of surfaces (29, 30, 42). Transcript levels for each the *fimABCD* operon genes were reduced in the Δ*gumB* mutant compared to the wild type by 15-85 fold in the RNA-Seq analysis. A reduction in *fimA* expression was similarly measured by Nanostring (59.2-fold) (Table 1) and by qRT-PCR (23.6-fold) (Figure 6A).

**Figure 6.**
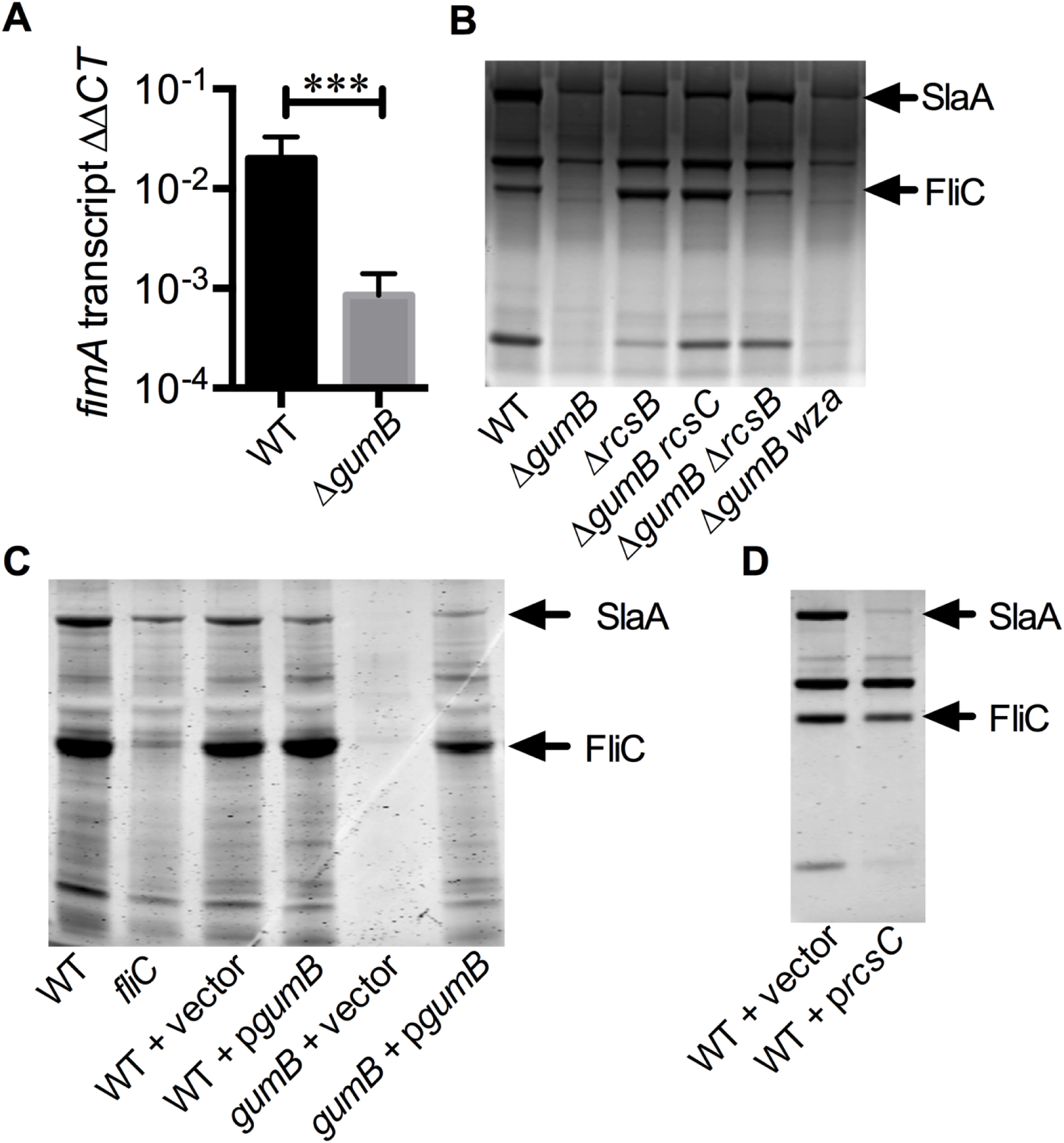
*S.marcescens* surface proteins including type I pilus biosynthesis (FimA) are decreased in the Δ*gumB* mutant and this is suppressed by inactivation of the Res system or multicopy expression of the *rcsA* gene. **A.** qRT-PCR of gene expression as noted above from cultures harvested at OD_600_= 1 in LB medium. p<0.001. n≥8. **B-D.** Representative image of Coomassie stained PAGE gels with sheared surface proteins from normalized cultures that were grown for 24 h at 30°C in LB medium. Vector control is pMQ132. p*gumB* is pMQ480; *prcsC* is pMQ615.

Surface protein fractions separated on PAGE gels revealed a reproducible and striking reduction in protein content (Figure 6B) that could be complemented by the wild-type *gumB* gene on a plasmid and (Figure 6C). The flagellin protein (FliC) was absent, as expected based on phenotypic and transcriptional analysis (14), and the surface-layer protein, SlaA, was reduced (Figure 6B-D). These protein bands have previously been extracted from gels and identified by mass spectrometry from similar preparations from this strain (40, 43). Similar to the Δ*gumB* mutant, activation of the Rcs system through multicopy expression of *rcsC* reduced levels of several surface proteins (Figure 6D), together the Δ*gumB* mutant had altered surface protein profiles.

Yeast agglutinations assay serves as a functional assay for type I pili levels from *S. marcescens* (30, 44). The time it takes to establish agglutination is inversely proportional to the relative fimbria level for a given genotype of *S. marcescens*. Whereas the wild-type bacteria caused yeast agglutination in less than 20 seconds, a no bacteria control and a pilus negative mutant (*fimC*) did not cause observable agglutination within one minute (Figure 7). The Δ*gumB* mutant was equally defective as fimbriae defective mutants, *fimA* and *fimC*, for yeast agglutination (Figure 7A, C). Agglutination could be restored to the Δ*gumB* mutant by the wild-type Δ*gumB* gene on a plasmid (Figure 7B) further indicating that GumB is necessary for pilus production.

**Figure 7.**
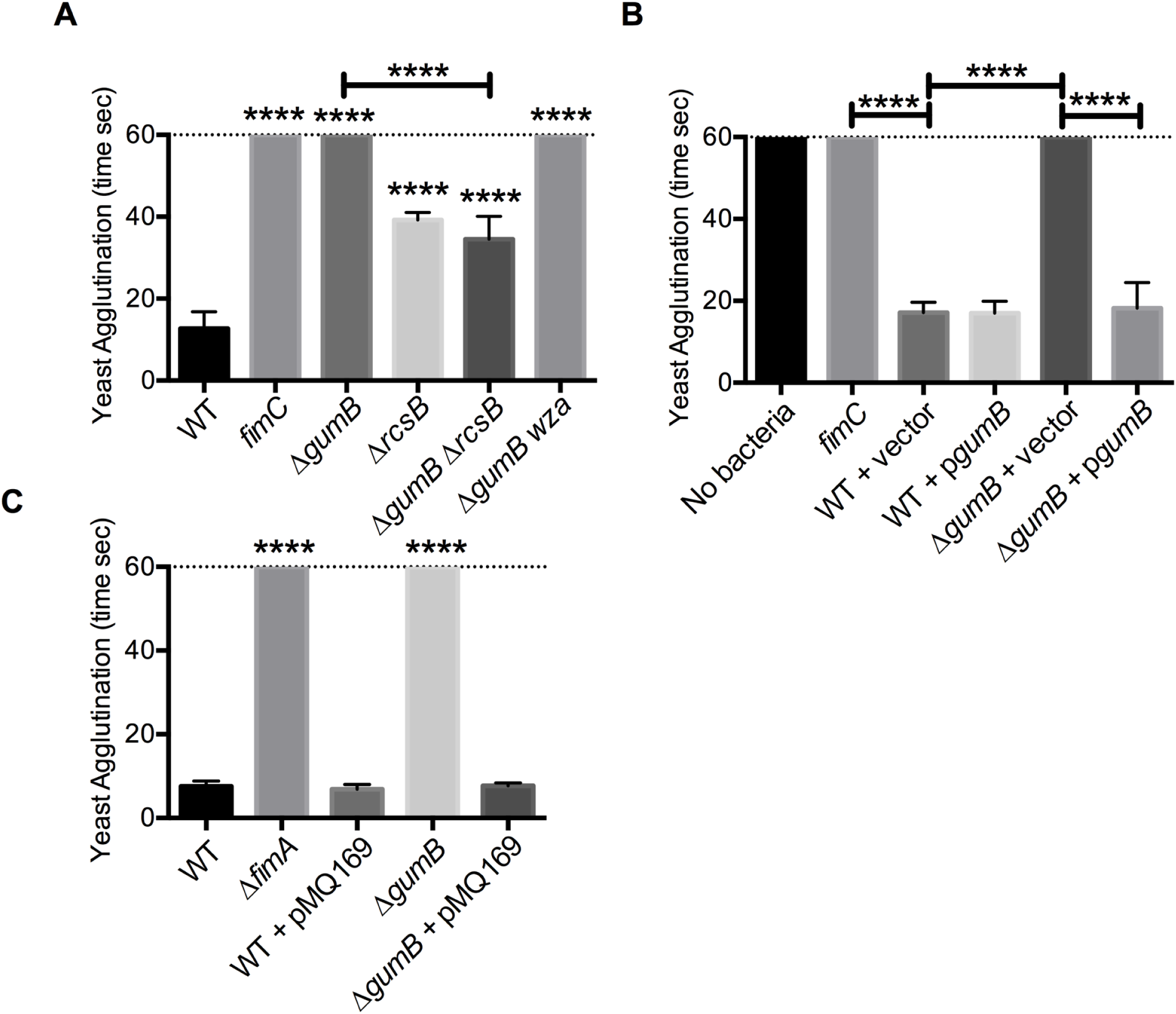
*S. marcescens* pilus production is altered in *gumB* and *rcsB* mutants. **A-C.** Timed pilus-dependent yeast agglutination indicates manose-dependent pilus activity. Faster agglutination time corresponds to more pili. Experiments were stopped at 60 seconds. Means and SD are shown, n≥3. Asterisks above the bar indicate significant difference from *WT,* p<0.0001. Vector control is pMQ132; *pgumB* is pMQ480. pMQ169 integration places the *fimABCD* operon under control of P_*lac*_

Because the GumB mutant had increased Rcs system activity that correlates with reduced FimA production, we tested whether inactivation of the Rcs system in the Δ*gumB* mutant could suppress the phenotype. The Δ*gumB* Δ*rcsB* double mutant could agglutinate yeast at levels similar to the Δ*rcsB* mutant, which was significantly more than the Δ*gumB* mutant (Figure 7A).

To further test whether the lack of fimbriae was solely responsible for the Δ*gumB* yeast agglutination defect, the chromosomal *fimABCD* operon was placed under control of a constitutive promoter by integration of plasmid pMQ169. This plasmid was previously used to demonstrate that a lack of fimbriae production was responsible for the *oxyR* mutant biofilm defect (30). The wild type with and without pMQ169 had similar yeast agglutination levels suggesting that the P*lac* promoter is of similar strength to the native promoter under the tested conditions (Figure 7C). The Δ*gumB* mutant with pMQ169 regained the ability to agglutinate yeast, indicating the fimbriae production was restored and was responsible for the yeast agglutination defect (Figure 7C).

### Proteomic analysis of surface proteins demonstrate differences between the wild type and Δ*gumB* mutant

To confirm the PAGE analysis and yeast agglutination data, sheared protein fractions from wild type and Δ*gumB* mutant were analyzed by mass spectrometry (Figure 8). Over 300 proteins were identified in these fractions, and while there was a significant correlation among expression of proteins found in both the wild type and Δ*gumB* mutant fractions (p<0.0001) (Figure S2), 22 proteins were measured at statistically different levels and several others were found only in one strain or the other, but were not significantly different (Figure 8). Of the 22, twelve were found in both strains, nine were measured from the wild-type strain only, and one was found only in the Δ*gumB* protein fractions. These include proteins relevant for phenotypes evaluated in this study. For example, the type I pilus protein FimA was found only in the wild-type samples (p=0.0032). Although not expected to be surface associated, some of the metalloproteases were differentially expressed. The SlpB protease was not found in the Δ*gumB* mutant fractions (p=0.0005); the PrtS protein was also found 1.8-fold higher in the wild-type samples; however, this did not reach significance (p=0.1375).

**Figure 8.**
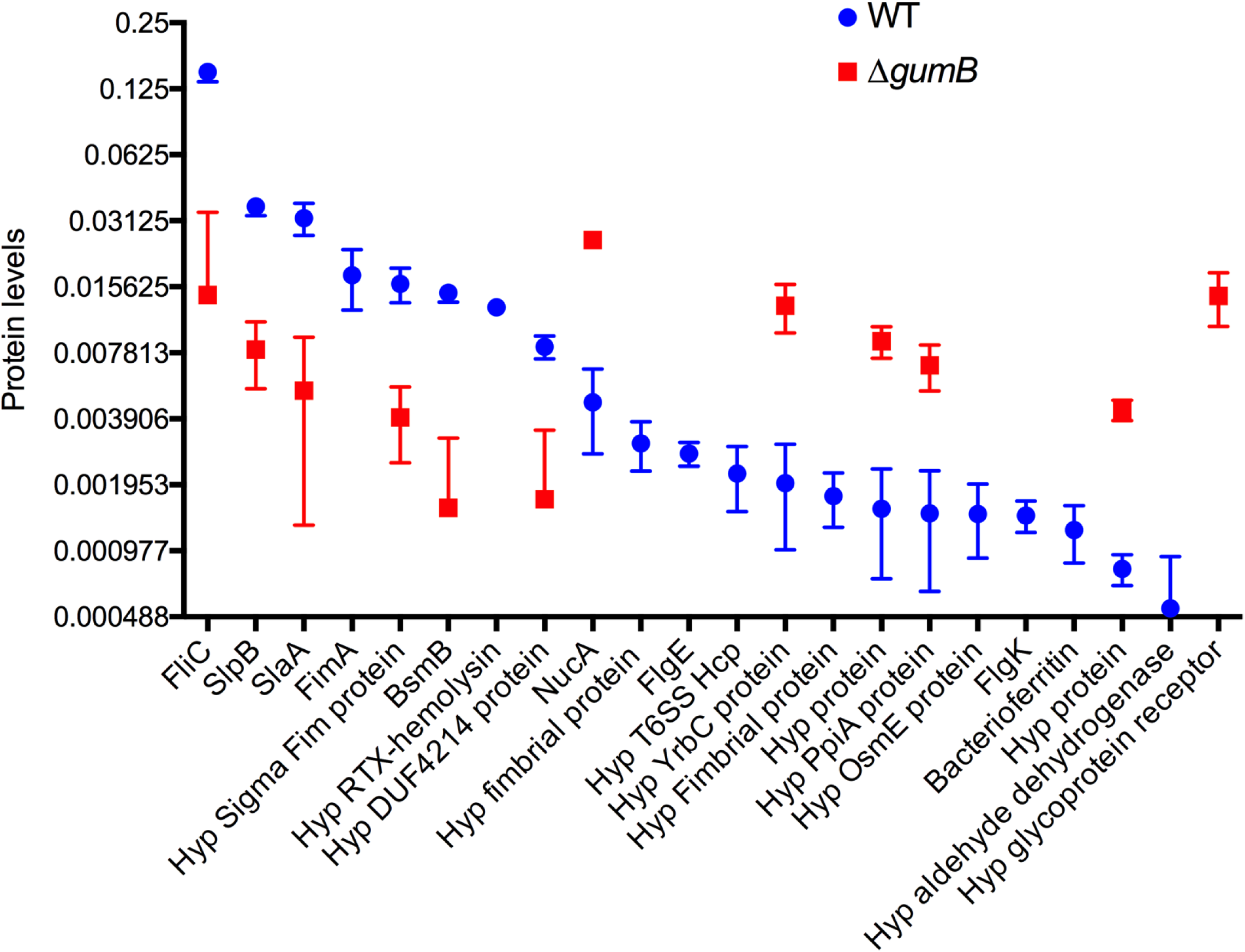
Proteomic analysis of surface protein fractions of the *WT* andΔ*gumB* mutant. Mean and SD of normalized spectral abundance factor (NSAF) are shown for each protein, n=3. Proteins data points not shown were undetected. All groups are 2-fold and significantly different between groups, *p<0 .05*.

Consistent with PAGE analysis (Figures 6B-D), a greater than 10-fold reduction in SlaA and FliC proteins were measured in the Δ*gumB* mutant compared to wild type (Figure 8, p<0.05). It is notable that the surface associated cytolysin ShlA was found in the wild type, but not the Δ*gumB* mutant, but this was not a significant difference (p=0.0827), likely due to the small sample size (n=3).

PSORTb V3 (45) and Bastionhub (46) were used to further characterize the surface faction proteins with respect to their predicted subcellular localization and secretion system profile (Tables 3-5). With PSORTb, of the 22 proteins with statistical differences, 2 were predicted to by cytoplasmic, 4 as not cytoplasmic, 2 as periplasmic, 11 as extracellular (such as FimA and flagellar proteins), 1 as outermembrane or extracellular, and 2 were undetermined. This suggests, that while fractions contained some cytoplasmic contamination, it was largely composed of extracytoplasmic proteins. Bastionhub software predicted 19 of the 22 proteins are secreted through Type I, II, III, IV, and VI secretion systems; however, *S. marcescens* strains do not generally have a type III secretion system.

**Table 3.** Significantly different expressed surface-associated proteins found in both strains.

| ORF<sup>a</sup> | Predicted Protein | WT/$\Delta gumB$<sup>b</sup> | P-value<sup>c</sup> | PSortB<sup>d</sup> | Predicted Secretion system<sup>e</sup> |
| --- | --- | --- | --- | --- | --- |
| SMDB11_2186 | FliC, flagellin | 10.39 | 0.001 | EC | II |
| SMDB11_0227 | FlgK, flagellar hook protein | 9.58 | 0.006 | NC | IV |
| SMDB11_0863 | Surface Layer Protein SlaA | 6.13 | <0.001 | EC | II |
| SMATCC274_R S07070 | DUF4214 domain-containing protein | 4.96 | 0.031 | EC/OM | I |
| SMDB11_1367 | SlpB metalloprotease | 4.48 | 0.007 | EC | I |
| SMDB11_1378 | BsmB | 4.07 | 0.005 | EC | I |
| SMDB11_2345 | Phenylacetic acid degradation protein | 2.98 | 0.035 | C | None |
| SMDB11_3835 | Peptidyl-prolyl cis-trans isomerase PpiA | -4.73 | 0.008 | P | II |
| SMDB11_1628 | hypothetical protein | -5.31 | 0.003 | NC | None |
| SMDB11_1061 | Nuclease, NucA | -5.49 | 0.005 | EC | II |
| SMDB11_3581 | hypothetical protein | -5.80 | 0.011 | NC | III |
| SMDB11_3593 | YrbC | -6.44 | 0.046 | NC | None |

<sup>a</sup> ORF number for reference strain Db11 (77) or strains ATCC274 (78) or WW4 (79) if the gene is not present in Db11.
<sup>b</sup> Ratio of Normalized SAF from WT compared to the $\Delta gumB$ mutant
<sup>c</sup> Student's T-test
<sup>d</sup> PSortB (45) subcellular localization analysis: C – cytoplasmic, EC – extracellular, NC – not cytoplasmic, OM – outer membrane, P – periplasmic, U – undetermined
<sup>e</sup> BastonHub secretion system prediction (46)

**Table 4.** Surface proteins only isolated from the wild type.

| ORF<sup>A</sup> | Predicted Protein | WT Avg NSAF<sup>B</sup> | SD | PsorB<sup>C</sup> | Secretion system<sup>D</sup> |
| --- | --- | --- | --- | --- | --- |
| SMDB11_3916 | FimA type I pilus subunit | 0.0176 | 0.0054 | EC | II |
| SMWW4_vlc32310 | RTX N-terminal domain | 0.0126 | 0.0008 | U | II |
| SMWW4_vlc31330 | Type-1 fimbrial protein | 0.0030 | 0.0008 | EC | I |
| SMDB11_3915 | PapA pili protein | 0.0028 | 0.0025 | EC | I |
| SMDB11_2216 | Flagellar hook protein FlgE | 0.0027 | 0.0003 | EC | II |
| SMWW4_vlc31580 | Hcp1, T6SS | 0.0022 | 0.0007 | EC | VI |
| SMDB11_0909 | Type-1 fimbrial protein | 0.0017 | 0.0005 | EC | II |
| SMDB11_2655 | Osmotically inducible lipoprotein E | 0.0014 | 0.0005 | P | II |
| SMDB11_2209 | Flagellar hook-associated protein FlgK | 0.0014 | 0.0002 | EC | II |
| SMDB11_2187 | Flagellar hook-associated protein FliD | 0.0013 | 0.0007 | EC | I |
| SMDB11_3798 | Bacterioferritin | 0.0012 | 0.0004 | C | None |
| SMDB11_1975 | DNA-binding protein H-NS | 0.0012 | 0.0021 | C | None |
| SMDB11_1941 | Outer membrane protein W | 0.0011 | 0.0008 | OM | II |
| SMDB11_3737 | ShlA | 0.0009 | 0.0010 | OM | I |
| SMDB11_3939 | Universal stress protein A | 0.0009 | 0.0008 | C | None |

<sup>A</sup> ORF number for reference strain Db11 (77) or strains ATCC274 (78) or WW4 (79) if
the gene is not present in Db11.
<sup>B</sup> Student's T-test
<sup>C</sup> PSorB (45) subcellular localization... C – cytoplasmic, EC – extracellular, NC – not
cytoplasmic, OM – outer membrane, P – periplasmic, U – undetermined
<sup>D</sup> BastonHub (46) secretion system prediction

**Table 5.**
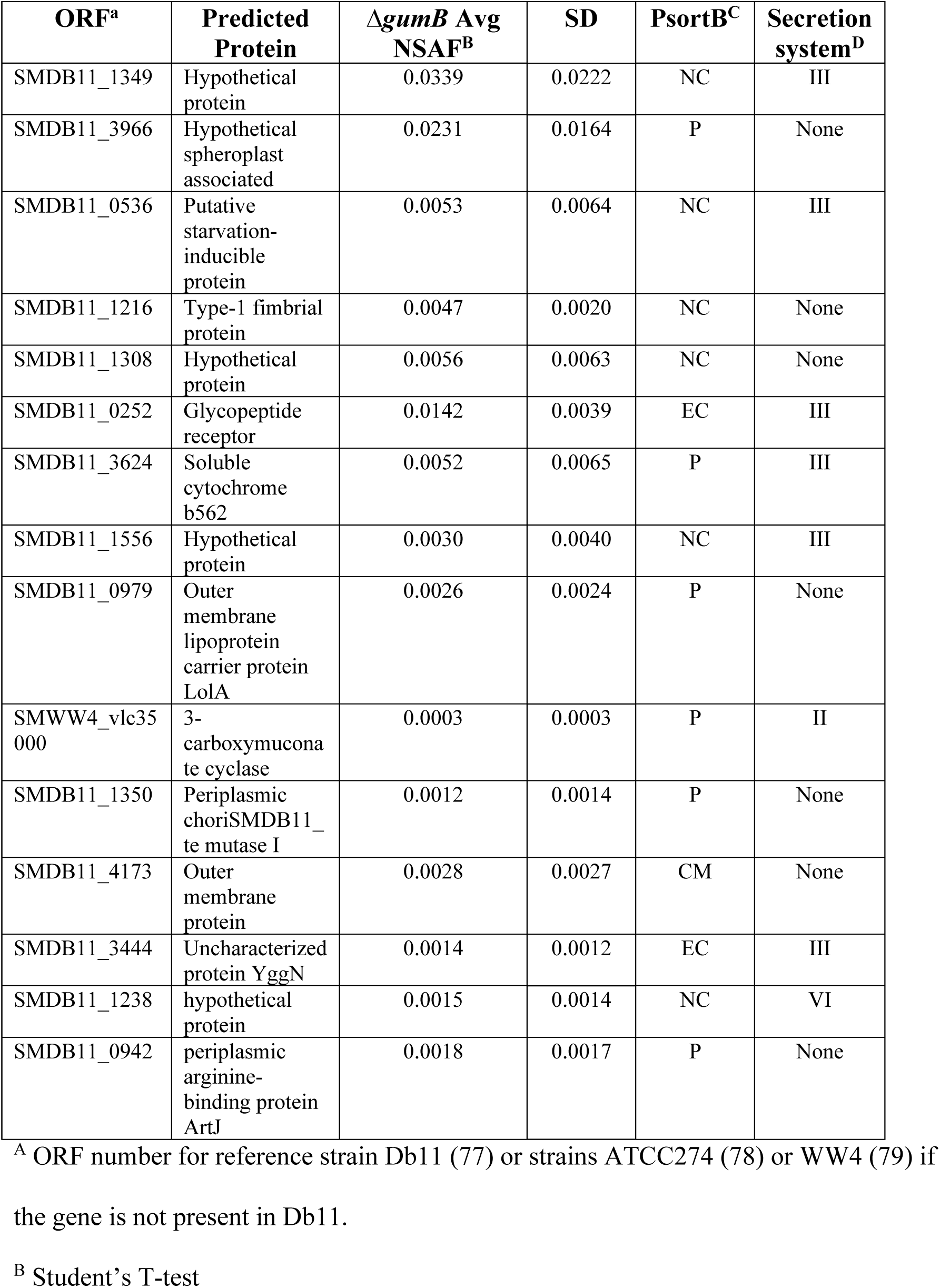
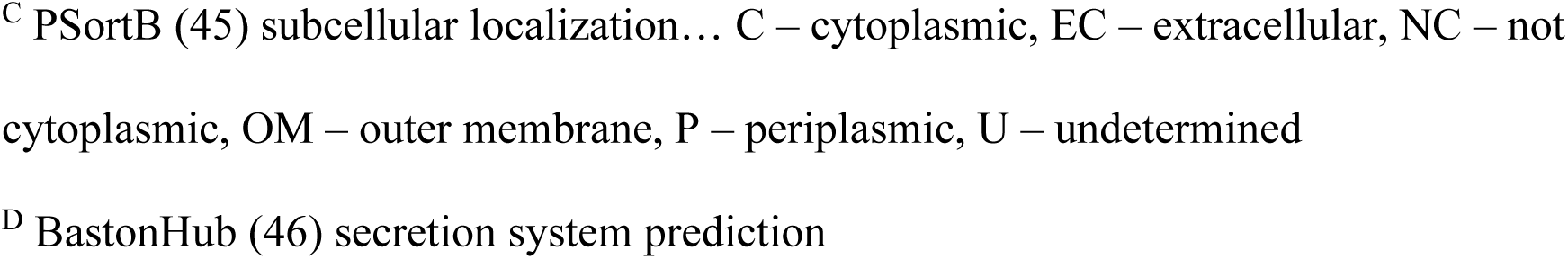
Surface proteins only isolated from the Δ*gumB* mutant.

## Discussion

This study investigated the impact of IgaA protein, GumB, on the global transcriptome and surface proteome of *S. marcescens*. The Δ*gumB* mutation was used as a tool to activate the Rcs system, a major envelope stress regulator, because of its key role in regulating virulence. Beyond this practical use of the *gumB* mutations in studying the Rcs system, natural mutations in *igaA* orthologs have been found in bacteria that were exposed to immune cells or the immune system *in vitro* and *in vivo*, so analysis of *igaA* mutants may be relevant to learn about bacteria exposed to immune system mediated stresses or isolated from chronic infections (47, 48).

A notable finding from this study was that many of the known virulence and hemolytic factors of *S. marcescens* were down regulated with Rcs activation. These include the *shlBA* cytolysin operon and its regulator *flhDC*, the phospholipase operon, the biosurfactant serratamolide (serrawettin W1), and metalloproteases genes. Consistently, secreted components of the Δ*gumB* mutant were significantly reduced in cytotoxic potency against a human corneal epithelial cell line. Induced expression of a metalloprotease gene, serralysin (*prtS*), was the only tested factor sufficient to restore cytotoxicity to the culture supernatants. These metalloproteases are well known cytotoxic factors from *S. marcescens* (31, 38, 39) and serralysin orthologs are found in many human pathogens including *Pseudomonas aeruginosa*. In fact, the *P. aeruginosa* alkaline protease, AprA, is closer in sequence to *S. marcescens* SlpB than SlpB is to the other *S. marcescens* metalloproteases (39). The reason that several bacterial species have genes for multiple serralysin proteases is not clear as these proteins have promiscuous protease activity and overlapping cytotoxic function (31). However, it has been hypothesized that they provide selective survival advantages during exposure to predatory bacteria present in soil and water (40). Beyond cytotoxicity and promoting survival versus predation, serralysin proteases are thought to influence the immune system through cleavage of immune modulators like cytokines and surface receptors (49, 50). For example, pretreatment of lungs with serralysin, remarkably increased influenza replication and pathology, indicating that serralysin metalloproteases can influence host-pathogen interactions (51).

Here we found a reduction of protease activity in the extracellular fractions of the Δ*gumB* mutant correlated with reduced transcription of three serralysin protease genes, *prtS*, *slpB*, and *slpE*. These results were similar to those found with mutation of the two-component regulator *eepR* (31, 43). However, in this study we observed no significant change in *eepR* or *eepS* levels in the Δ*gumB* mutant versus the wild type, suggesting that the role of Rcs in control of these metalloprotease genes is independent of EepR. Beyond *S. marcescens*, a role for the Rcs system in controlling metalloprotease production was suggested in a RNA-seq study with *Proteus mirabilis*, where the genes for the IgA-degrading ZapA and ZapB metalloproteases, was the most highly upregulated gene in bacteria with an inactivated Rcs system (*rcsB* mutation) compared to the wild type (52). This suggests a conserved role for the Rcs system in repressing this group of important proteases.

Beyond toxin and protease downregulation, several putative fimbrial/pili genes were observed to be altered in expression including the *fimABCD* operon which codes for a mannose dependent type I pilus, a major biofilm factor for *S. marcescens* (29, 30, 42, 44). Data presented here suggested that the FimA protein was Rcs regulated and had reduced expression in the Δ*gumB* mutant. This was independent of the capsular polysaccharide, as mutation of an essential gene in capsular polysaccharide biosynthesis did not affect the FimA levels or FimA-dependent phenotypes in the Δ*gumB* mutant background. Interestingly, while the Δ*gumB* mutant phenotypes indicated that hyper-activity of the Rcs system leads to an almost complete loss of FimA production, the inactivation of the Rcs system through mutation of *rcsB* also reduced FimA production to intermediate levels. Similar effects were seen with the Δ*gumB* Δ*rcsB* and Δ*gumB rcsC* double mutants. Together, these suggest that too much or too little Rcs activity reduce bacterial adhesion production. It is likely that a number of signals, such as interaction with surfaces influence Rcs control over adhesion development, and this in turn likely impacts biofilm formation. Temporal and spatial control of the Rcs system during biofilm formation has been reviewed by the Clarke group (1, 53).

A role for the Rcs system in control of fimbriae has been suggested for other organisms such as *E. coli* (54, 55), *K. pneumoniae* (56), *P. mirabilis* (52), and *S. enterica* (27). The Rcs system appears to inhibit fimbriae/pilus gene transcription in all but *P. mirabilis*, where it has a positive regulatory role. Consistently, an *rcsB* mutant of *P. mirabilis* was highly defective in biofilm formation on abiotic surfaces at early (8 h) and intermediate (24 h) time points, but caught up by 48 h. These results are consistent with the lack of an adhesin, and correlated with down regulation of the *mrpA* and *pmfA* fimbrial genes (52). Other studies have focused on the positive role of the Rcs system in biofilm formation through regulation of extracellular polysaccharides (15, 27, 57).

Thus, a major outcome of this study is the extent to which the *S. marcescens* GumB protein controls known adhesive factors beyond the capsular polysaccharide. In a study with *S. enterica*, it was predicted that IgaA proteins enacts fine tuning of the sessile existence within biofilms through repression of flagella and increased expression of extracellular polysaccharides (27).

While this study provides supportive data for Rcs regulation of primary attachment factors like fimbrial adhesins, defining the role of the Rcs system in early and later stages of biofilm formation by *S. marcescens* will require further study. The results of this study suggest that Rcs regulated genes likely influence multiple stages of the biofilm life cycle. For example, the *bsmB* gene and BsmB protein were found at reduced levels in the Δ*gumB* mutant in this study. BsmB was found by Labbate and colleagues to be quorum sensing regulated and to fine tune biofilm formation in *S. marcescens* strain MG1 (42, 58). In contrast to what is observed in *Pseudomonas* species (44, 59), the flagella of *S. marcescens* does not appear to be important for early stages of biofilm formation, although they could certainly play a major role in later stages of biofilm formation including dispersal or distribution of biofilms in more complex environments (44, 59). The reduced expression of PpiA, a peptidyl-prolyl cis-transferase, which had lower gene expression in the Δ*gumB* mutant, may affect fimbria production; related peptidyl-prolyl cis-transferases SurA of *E. coli* and FkpA of *S. marcescens* are necessary for pilus production and biofilm formation (44, 60). Surprisingly, the *S. marcescens* serralysin proteins can also contribute to biofilm formation, with expression of PrtS (also known as PrtA) being required for full levels of biofilm formation through an as yet unknown mechanism (61). As the Rcs system senses interaction of bacteria with surfaces (54), it is likely that Rcs mediated control of these multiple factors has a profound effect on several stages of biofilm formation. Numerous other genes were predicted to be involved in survival were upregulated in the Δ*gumB* mutant. These include capsule biosynthesis genes that have been associated with pathogenesis (62, 63), uncharacterized genes similar to osmolarity induced genes like *osmB,* and a predicted metallo-beta-lactamase gene which likely affect stress and antibiotic tolerance.

The transcriptome of *S. marcescens* was notably changed with the activation of the Rcs system (Δ*gumB* mutant). Similarities have been revealed with transcriptomic studies using mutations that inactivate Rcs system in *E. coli* (54, 55), *Erwinia amylovora* (64), *Klebsiella pneumoniae* (56), *P. mirabilis* (52), *S. enterica* (26, 27), *S. marcescens* (15), and *Yersinia entercolitica, pestis, and pseudotuberculosis* (57, 65, 66). However, with the exception of two microarray studies with *S. enterica* serovar Typhimurium (26, 27), previous studies transcriptomic studies of the Rcs system have used *rcsB*, *rcsC* or *rcsD* mutants to inactivate the system rather mutations that induce the Rcs response. This may be due to the fact that *igaA* is an essential gene in *E. coli* and *S. enterica*, and likely other organisms (5) (8, 9). Therefore, the *Salmonella* studies used strains with point mutations in *igaA* that create R188H or T191P changes and activate the Rcs system (26, 27), whereas our study used a full deletion of the *gumB* open reading frame. The two *Salmonella* studies varied with the extent of the impact of *igaA* mutation on the transcriptome, one concluded that almost 20% of genes had differential expression and the other 2%. These are reasonably similar to the ∼15% for *S. marcescens* in this study. Commonalities include increased expression of capsular polysaccharide, and osmotic protection genes, and decreased expression of flagellar and predicted virulence genes. An exception being that the flagellar *fliPQR* operon was highly upregulated in the *igaA* mutant of *S. enterica*. With *S. marcescens*, we observed a down regulation of ∼20-36 fold for *fliPQR* (26, 27). It was suggested that RcsA is necessary for *fliPQR* expression in *S. enterica*. In one study *rcsA* expression was >12-fold higher in the *igaA* mutant (26, 27). By contrast, in *S. marcescens ΔgumB*, *rcsA* expression was down ∼1.5 to 2-fold, which may account for some or all of the difference between species with respect to *fliPQR* expression. Other similarities include the general trend for decreased expression of virulence genes with Rcs activation, even though there are many different virulence mechanisms between the species. This outcome correlates with studies that have demonstrated reduced virulence phenotypes of *igaA* and *gumB* mutants (7, 11, 13). Additional differences in between the species include increased expression of *S. marcescens* specific genes for the nuclease, *nucA*, and chitinases associate genes in *S. marcescens*. Among these were the chitin binding protein gene, *cpb21*, which is similar to a *Vibrio cholera* protein involved in biofilm formation (67), and various chitinases that were upregulated in the Δ*gumB* mutant. It was also noted that surface layer protein A (SlaA) was reduced in the Δ*gumB* mutant by proteomics. Although this gene did not have significantly reduced transcription in the Δ*gumB* mutant, it was nevertheless decreased in the PAGE and proteomic analyses. This may be due to the observation that genes for the type 1 secretion system (T1SS) components *lipC* and *lipD*, which happen to be adjacent genes and are required for SlaA secretion (68), have significantly reduced expression levels in the *gumB* mutant. Although the surface layer function in *S. marcescens* is not well understood. The S-layer of other ocular pathogens contributes to their virulence and host-pathogen interactions in animal models (69, 70). Reduced expression of T1SS proteins was also reported for *rcsB* mutants of *P. mirabilis* (52).

The *in vitro* transcriptomic analysis was largely in agreement between the Nanostring and RNA-Seq approaches and these supported previous findings regarding transcriptional changes in the Δ*gumB* mutant (13, 14). Regarding the *in vivo* versus *in vitro* analysis, it was clear that many of the tested genes were expressed at relatively higher levels in the rabbit eye including virulence genes. Notably, however, there were several variables to this analysis that must be considered such as growth temperature, 30°C in vitro versus 32-33°C for the rabbit cornea, and potential differences in growth phase, OD_600_ = 1 versus 24 post-inoculation. The Rcs system was expected to be activated more highly in the *in vivo* samples given the hostile nature of the ocular surface (24). Consistent with the Rcs activation of the Δ*gumB* mutant were lower levels of flagellar and pilus gene production *in vivo*, but contrasted with increased levels of *shlBA* and *slpE*. This may be due to activation of these genes by other transcriptional regulators in the eye such as FlhDC (17, 37) or EepR (31, 43).

In conclusion, this study adds to our understanding of an IgaA ortholog’s global impact on bacterial transcriptomes and surface proteins. This revealed changes that manifested in clear loss of extracellular cytotoxic protease production and attachment associated fimbrial/pilus proteins. Data also suggests that GumB is a major transducer of signals that impact early and later stages of biofilm formation, the focus of subsequent research. Results from this study will help direct future mechanistic studies to understand the role of the Rcs system in controlling microbial pathogenesis.

## MATERIALS AND METHODS

### Strains and media

Strains of *S. marcescens* were all derived from keratitis isolate K904 (71) and are listed in Table S1. Bacteria were grown in lysogeny broth (LB) or on LB agar (72). Kanamycin was used at 100 µg/ml, gentamicin at 10 µg/ml, and tetracycline at 10 µg/ml. Bacteria were grown at 30°C except where noted.

### Mutagenesis

To make the *fimA* deletion strain, lambda red recombineering was performed using pMQ538 as previously described (16). The entire open reading frame except the first four codons were replaced by the kanamycin resistance cassette from pKD4 (73). Primers to amplify to perform this were 5239 and 5240. Primers to verify the insertion were 1879 and 5241. Primer sequences are listed in Table S1.

### RNA analysis and RNA-Seq

Cultures were grown to OD_600_=0.5, then subcultured 1:10 in 5 ml of LB medium, and allowed to grow until OD_600_=1.0. An aliquot (0.4 ml) of each culture was directly added to RNA-protect (Qiagen), processed according to the manufactures specifications including DNase I treatment as previously described (74)), and frozen at −80°C. qRT-PCR was performed as previously described (74). Transcripts were normalized using 16S expression; primers are listed in Table S1.

For RNA-sequencing analysis (RNA-Seq), strain K904 and K904 Δ*gumB* were streaked to single colonies on LB agar plates, incubated for 20 hours at 30°C, and single colonies were used to inoculate 5 ml of LB medium. Three tubes were used per strain, and in all cases tubes were aerated on a tissue culture rotor. Cultures were grown to OD_600_=0.5, then subcultured 1:10 in 5 ml of LB medium, and allowed to grow until OD_600_=1.0. An aliquot (0.4 ml) of each culture was directly added to RNA-protect (Qiagen), processed according to the manufactures specifications including DNase I treatment as previously described (74), and frozen at −80°C. This was repeated every day for three days with identical timing. The three RNA samples from each of the three days were pooled into one sample for each day. RNA was analyzed for chromosomal DNA contamination using primers for 16S rDNA, and none was observed when amplified for 28 cycles followed by analysis on an agarose gel. Total RNA was extracted as above, and samples were sent to GeneWiz, Inc. (New Jersey) for processing, including depletion of ribosomal RNA, RNA library preparation, sequencing with an Illumina HiSeq2500 platform in a 2×150 bp single-read configuration. Gene hit counts were determined using featureCounts (Sunbread). DESeq2 was used for differential gene expression analysis and the Wald test was used to generate p-values and log2 fold changes. KEGG gene ontology was performed on differentially expressed genes (2-fold difference from the wild type, p<0.05) using GhostKOALA (28).

RNA for the nanoString nCounter platform was harvested from corneas following the protocol of Xu, et al. (75) with a few exceptions. Corneas were removed with a trephine and immediately frozen in liquid nitrogen and stored at −80°C until RNA was harvested. Corneas were homogenized as described above. Following phenol chloroform extraction and ethanol precipitation, RNA was run through a ZymoResearch RNA concentrator column, adjusted to 800 ng/µl and sent to the University of Pittsburgh Genomics Research Core for nCounter analysis. Results were analyzed using nSolver software and results were uploaded to Gene Expression Omnibus (accession number GSE155486). Probes with counts equal or lower than the geometric mean of the negative control probes were considered below the limit of detection and omitted, such that the number per group varied from 3-5 for the in vivo grouping and 5-7 and genes with less than 2 samples were omitted from analysis. Two-tailed Student’s T-tests were performed on Log2 transformed data.

### Rabbit keratitis model

This study conformed to the ARVO Statement on the Use of Animals in Ophthalmic and Vision Research and was approved by the University of Pittsburgh Institutional Animal Care and Use Committee (IACUC Protocol 16098925). As previously described (11), bacteria were grown overnight, adjusted to ∼500 CFU in 25 µl, and injected intrastromally into the right corneas of New Zealand white rabbits. At 24 h post-injection, animals were euthanized with an intravenous overdose of Euthasol solution following systemic anesthesia with 40 mg/kg ketamine and 4 mg/kg of xylazine. The corneas were removed with a 10 mm trephine and immediately frozen in liquid nitrogen and stored at −80°C until RNA was harvested. Corneas were homogenized with an MP Fast Prep-24 homogenizer using lysing matrix A tubes (MP Biomedicals). Flowing phenol chloroform extraction and ethanol precipitation, RNA was run through a ZymoResearch RNA concentrator column, adjusted to 800 ng/µl and sent to the University of Pittsburgh Genomics Research Core for nCounter analysis. RNA for the nanoString nCounter platform was harvested from corneas following the protocol of Xu, et al. (75).

### Proteomics and protein analysis

Surface protein fractions were obtained as previously described (76). Briefly, cultures were grown overnight (18-20 h) in LB broth and adjusted to OD_600_ = 5.0. An aliquot (1.5 ml) was centrifuged (13,000 x g for 1.5 min), washed with PBS (0.5 ml), and suspended in PBS (1 ml) in a microfuge tube. The tube was vortexed (Vortex Genie) for 2 min on the maximum setting, 0.5 ml of cold PBS was added and the tube was centrifuged at 21,000 x g for 1 min, and the supernatant was passed though a 0.22 µm PVDF filter. To an 800 µl aliquot of the filtrate, 200 µl of trichloroacetic acid (Sigma) was added and mixed by inversion (5x) and placed on ice for 30 min. The tubes were centrifuged at 21,000 x g for 15 minutes at 4°C, the supernatant was discarded, and the pellet was washed with acetone twice (400 and 200 µl, respectively). The pellets were dried at 65°C and stored at −20°C until being used for PAGE gels (12%), stained with Coomassie Brilliant Blue, and imaged with a Li-COR Odyssey imager, or sent to MS Bioworks (Ann Arbor MI) for proteomic analysis. PAGE gels were stained with Coomassie. Samples were washed with ammonium bicarbonate (25 mM) and acetonitrile, reduced with dithiothreitol (10 mM) at 60°C and alkylated with iodoacetamide (50 mM) at room temperature. The proteins were digested with trypsin for 4 h at 37°C, the reaction was stopped with formic acid. Samples were analyzed by nano-LC-MS/MS with a Waters M-class HPLC system with a Fusion Lumos (ThermoFisher). Peptides were separated on a trapping column and eluted over an analytical column (75 µm) at 350 nL/min, both of which with Luna C18 resin. The mass spectrometer was operated with the Orbitrap operating at 60,000 FWHM for MS and 15,000 FWHM for MS/MS. The instrument was run with a 3s cycle for MS and MS/MS and APD was enabled and 5 h of instrument time was used for each sample. Data was analyzed with Mascot software (Matrix Science) and parsed into Scaffold software. Data were filtered using 1% protein and peptide FDR and required at least two unique peptides per protein. Normalized SAF values were determined and compared between strains using a two-sided Student’s T-test.

### Yeast agglutination assay

Agglutination of *Saccharomyces cerevisiae* was performed as previously described (40, 44). Bacteria were grown overnight in LB broth, washed and adjusted to OD_600_=10.0 in PBS and mixed with dried yeast (Sigma YSC2) in PBS (0.02g/ml) and 25µl of bacterial suspension or negative control PBS alone and 25µl of yeast were combined on a microscope slide on a rotating platform shaker (Barnstead Multipurpose Rotator) set to half maximum speed. The time of agglutination was measured using a stopwatch, with a maximum time of 60 second.

### Statistical analysis

Other than noted above, one-way ANOVA with Tukey’s post-hoc test was used to compare multiple groups and two-tailed Student’s T-test or Mann-Whitney were used for pairwise comparisons. Spearman analysis was used for correlation analysis. Analysis was performed with Graph Pad Prism software.

### Data availability

Raw Transcriptomic data has been deposited at the accession number GSE155486 and GSE152115 (NanoString) and GSE151031 (RNA-Seq). Other data is available upon request.

## ACKNOWLEDGEMENTS

The authors acknowledge Kara Lehner and Jake Callaghan for aiding in cloning and strain building and Michael Ford for help with proteomic analysis. This work was supported by National Institutes of Health grants R01EY027331 (to R.S.) and CORE Grant P30 EY08098 to the Department of Ophthalmology, University of Pittsburgh School of Medicine, Pittsburgh, PA. The Eye and Ear Foundation of Pittsburgh and an unrestricted grant from Research to Prevent Blindness, New York, NY provided additional departmental funding.

